# Epigenetic insights into neuropsychiatric and cognitive symptoms in Parkinson’s disease: A DNA co-methylation network analysis

**DOI:** 10.1101/2023.07.20.549825

**Authors:** Joshua Harvey, Adam R. Smith, Luke S. Weymouth, Rebecca G. Smith, Isabel Castanho, Leon Hubbard, Byron Creese, Kate Bresner, Nigel Williams, Ehsan Pishva, Katie Lunnon

## Abstract

Parkinson’s disease is a highly heterogeneous disorder, encompassing a complex spectrum of clinical presentation including motor, sleep, cognitive and neuropsychiatric symptoms. We aimed to investigate genome-wide DNA methylation networks in post-mortem Parkinson’s disease brain samples and test for region-specific association with common neuropsychiatric and cognitive symptoms. Of traits tested, we identify a co-methylation module in the substantia nigra with significant correlation to depressive symptoms and with ontological enrichment for terms relevant to neuronal and synaptic processes. Notably, expression of the genes annotated to the methylation loci present within this module are found to be significantly enriched in neuronal subtypes within the substantia nigra. These findings highlight the potential involvement of neuronal-specific changes within the substantia nigra with regard to depressive symptoms in Parkinson’s disease.

## Introduction

Parkinson’s disease (PD) is the second most common neurodegenerative disease and is the fastest growing in prevalence of all neurological disorders, estimated to affect 6.1 million individuals worldwide based on a 2016 census^1^. Clinically, PD is defined by its cardinal motor symptoms (resting tremor, bradykinesia, rigidity and postural instability)^2^, but highly prevalent features of the disease encompass a range of cognitive and neuropsychiatric symptoms^3^. Common symptoms, reported in a high proportion of patients, include depression^4^, anxiety^5^, psychosis (most prominently hallucinations and delusions)^6^, apathy^7^, cognitive impairment and dementia^8^. The cumulative effect of these secondary symptoms greatly increases disease burden for patients and complicates treatment^9,10^. As examples, psychosis is an associated factor to increased nursing home placement^11^, mortality and caregiver burden in PD^12^. Dopaminergic therapies, highly prescribed for motor symptom treatment, reportedly increase individual risk for the emergence of psychosis symptoms^13^. The development of these secondary symptoms is not always timed after the diagnosis of the primary motor disorder, for example, depression is a common manifestation in premorbid PD and has been associated as a risk factor for motor symptom development^14–16^. Furthermore, the therapies that exist for these non-motor symptoms are currently minimally effective, despite the considerable disease burden they represent.

Although the occurrence of neuropsychiatric and cognitive symptoms in PD is much more common than in age-matched populations^9, 10^, individual to individual level susceptibility to these secondary features is highly variable^17, 18^. Genetic liability has been implicated, for example a recent genome-wide association study (GWAS) of cognitive progression in PD highlighted the contribution of risk genes such as *GBA* with worsening cognitive decline over time^19^ and meta analyses of the gene have shown an association to the emergence of psychosis and depression symptoms^20^. However, given the high levels of heterogeneity within the condition, PD secondary symptoms likely share a complex underlying etiology, owing to additional factors aside from genetics. One potential contributing factor is epigenetic changes, which play an intermediary role between genetic and environmental risk, and regulate gene expression^21^. DNA methylation, which refers to the reversible addition of methyl groups to cytosines typically in a CpG dinucleotide, is the most studied epigenetic mechanism in neurological disorders^22^. Indeed, several studies have shown robust alterations in DNA methylation in a number of genes in different neurodegenerative diseases, in both the brain and blood, including Alzheimer’s disease (AD)^23–25^, PD^26–28^ and Dementia with Lewy bodies (DLB)^29^. Interestingly, associations have also been reported for secondary symptoms of these neurodegenerative disease, for example with psychosis symptoms in AD^30^ or cognition in PD^27^. However the analysis of DNA methylation signatures in relation to PD secondary symptoms is understudied and has predominantly been undertaken in peripheral tissues such as blood^31^.

In the current study we investigated the relationship between DNA methylation patterns and the occurrence of key secondary symptoms in PD (dementia, hallucinations, depression, anxiety, aggression, sleep disorder), using weighted gene correlation network analysis (WGCNA) in multiple disease-relevant brain regions. Subsequently, gene ontology and cell type enrichment analysis were performed on the genes comprising the significant modules to identify dysfunctional pathways and the cell types likely driving this. We highlight a core finding of a co-methylation module specific to the substantia nigra, significantly correlated to depressive symptom presentation and significantly enriched for neuronally relevant synaptic terms. Assessing the expression of genes annotated to this module found enriched expression in specific neuronal sub-populations within the substantia nigra, indicative of neuronal changes within this region that may play a role in the development of depressive symptoms within PD.

## Results

### A cohort to assess DNA methylation signatures of PD neuropsychiatric and cognitive symptoms

Our study comprised a cohort of 97 idiopathic Parkinson’s Disease (PD) patients with post-mortem DNA methylomic profiling conducted on the Illumina Infinium 450K array (Figure 1A). Three brain regions were assessed: the substantia nigra (SN, n = 88), caudate nucleus (CN, n = 82) and prefrontal cortex (FC, n = 88), with the majority of cases having all brain regions represented in this dataset (Figure 1A,C). PD patients had a mean age of 78.25 years at death (SD = 6.17) with the average patient having had PD symptoms for over ten years (mean = 12.63, SD = 8.28). Pathologically, these patients predominantly showed late stage PD-associated Lewy body (LB) pathology^12^ (LB Braak stage mean = 5.54, SD = 0.81) with relatively mild AD-associated neurofibrillary tangle (NFT) pathology^32, 33^ (NFT Braak stage mean = 1.91, SD = 0.72) (Figure 1B).

**Figure 1:**
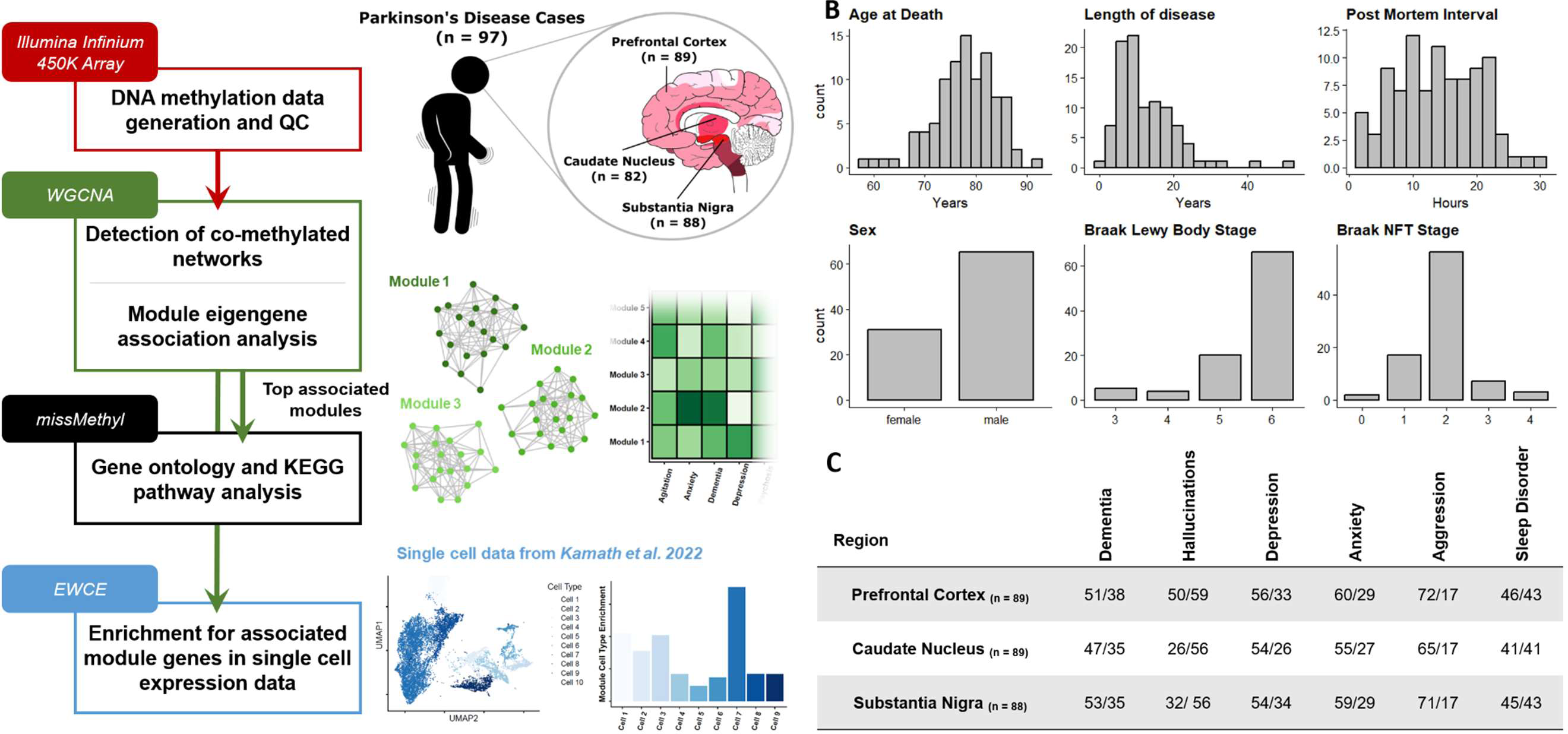
Overview of study design and samples used. **A)** Sample information and data analysis flowchart as detailed in Methods **B)** Demographic summaries of PD patients profiled with histograms or bar-charts representing overall numbers for each variable. Length of disease corresponds to time interval between recorded age of PD diagnosis and age at death. **C)** Summary table of sub-symptoms tested in primary WGCNA association analysis. Symptoms were annotated for binary status in the ante-mortem clinical records. Symptom prevalence per each brain region tested is shown and is annotated as Absent/Present.

The primary hypothesis of this study posits that the prevailing neuropsychiatric and cognitive manifestations observed in PD exhibit a distinctive epigenetic profile in the brain, distinguishing individuals presenting with these symptoms from those who do not. To test this hypothesis, we annotated binary symptom prevalence from antemortem clinical records for six phenotypes: dementia, hallucinations, depression, anxiety, aggression, and sleep disorder (Methods, Figure 1C). The majority of these sub-symptoms showed overlap in their presentation and demographic differences (Supplementary Figure 1, Supplementary Table 1) as could be expected with cumulative disease burden^34^. To identify DNA methylation signatures associated with the phenotypes of interest, we investigated co-methylation changes by implementing weighted gene correlation network analysis (WGCNA). This was also useful as a strategy to reduce the number of features and increase statistical power, given our modest sample size for conducting epigenome-wide association studies (EWAS), particularly when examining binary variables related to phenotypes of interest.

### DNA co-methylation networks show brain region specific correlation to depressive and aggression symptoms

To identify co-methylated modules within each brain region, we followed a standardized WGCNA protocol (Methods), and tested their association with sub-trait presentation, after regressing out key covariates (age, sex, technical batch, proportions of neurons, post-mortem interval (PMI)). The number of detected modules differed across each brain region, with 27 modules identified in the SN (Supplementary Figure 2), eight in the FC (Supplementary Figure 3) and 18 in the CN (Supplementary Figure 4). The correlation of these modules to trait presentation also differed across brain regions. Stronger module-trait correlations were observed in the SN and CN, with two modules passing the Bonferroni significance threshold for the number of tests within each trait association (Figure 2, SN: P < 0.0019, CN: P < 0.0028), with no significant correlations in the FC (Figure 2). The significant SN module correlated with depressive symptoms in PD (Spearman’s Coefficient = 0.33, P = 0.0016) and was comprised of 1,375 distinct methylated loci, whilst the significant CN module that was significantly correlated to aggression presentation (Spearman’s Coefficient = 0.35, P = 0.0015) was comprised of 475 distinct methylated loci.

**Figure 2:**
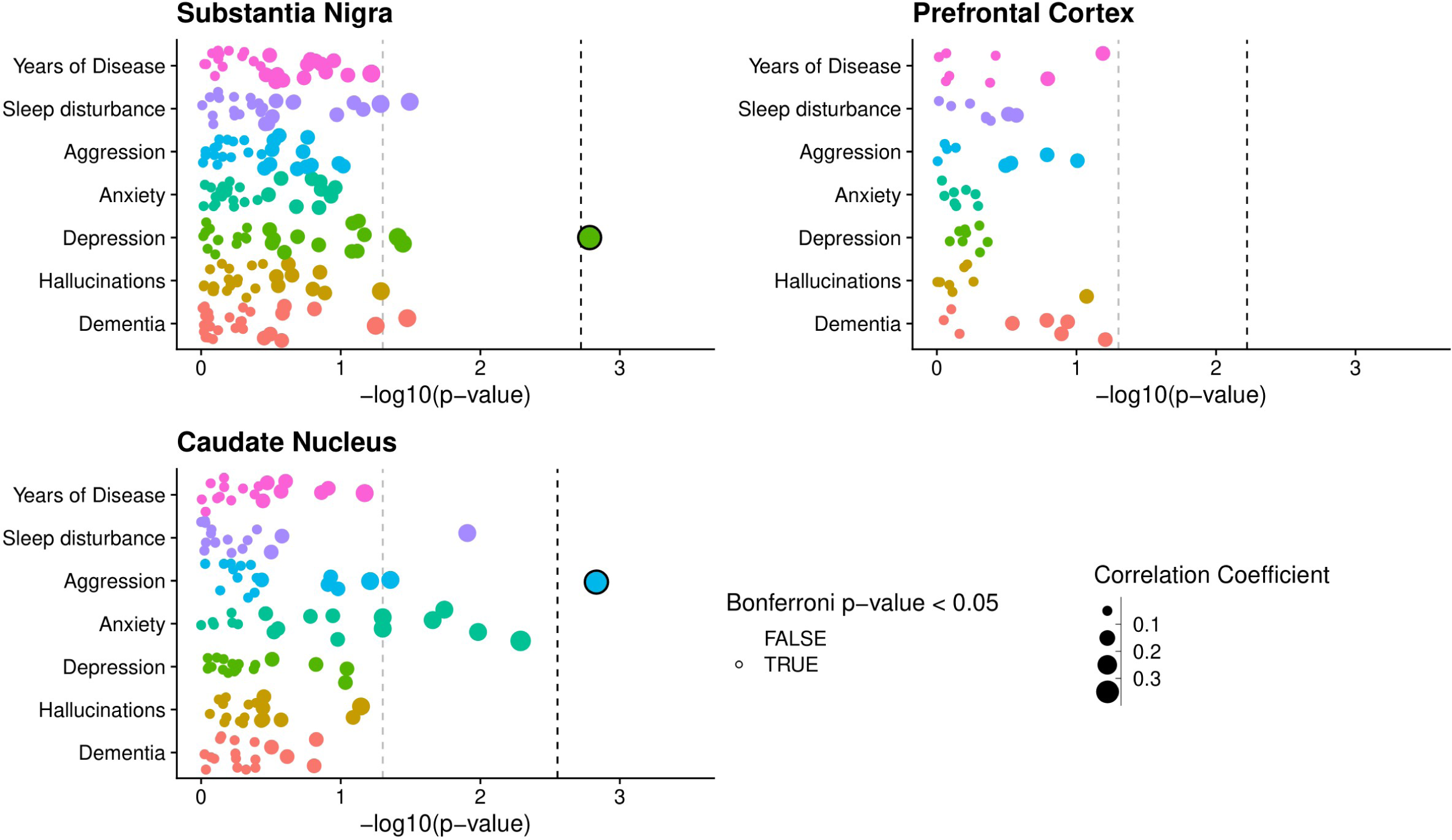
Co-methylation network association to sub symptom occurrence in PD. Points represent individual module eigengenes for the substantia nigra (n = 27), frontal cortex (n = 8) and caudate nucleus (n = 18), repeatedly tested using correlation analysis in association with traits displayed along the y axis. Points are colored by the trait they are being tested for association with and sized by the absolute correlation coefficient value of the association. For clinical binary traits Spearman’s correlation was used, whilst for years of disease Pearson’s correlation was used. The -log10(p-value) of the association tests is displayed along the X-axis. The gray dashed line represents P-value = 0.05, whilst the Black dashed line represents the Bonferroni correction threshold for each brain region, controlling for the number of tests within each trait association, equivalent to 0.05 divided by the number of module eigengenes per region (SN: P < 0.0019, CN: P < 0.0028, FC P < 0.006).

When assessing module membership of these two significant modules, a weak but significant correlation was observed between P-value significance of depressive symptom association and module membership for methylated loci within the SN depression associated module (Pearson’s Coefficient = 0.12, p-value = 1.24 x 10^-5^, Supplementary Figure 5A). By contrast the CN aggression associated module does not show any indication of correlation between module membership and probe significance from the aggression symptom association (Pearson’s Coefficient = 0.07, p-value = 0.13, Supplementary Figure 5B).

Although no further modules passed our threshold for multiple testing correction, several other modules in these regions did show nominal significance in their correlation with trait presentation (Figure 2, Supplementary Figure 2, Supplementary Figure 4). Of particular note, a set of four modules in the CN all showed correlation with anxiety symptoms. However, we have focused our downstream analyses on the Bonferroni-significant module identified in the SN (with respect to depression) and the CN (associated with aggression), henceforth referred to as the DepressionSN module and the AggressionCN module, respectively.

To test whether the association of the DepressionSN module was affected by onset of depression before motor symptoms we subset the depression group based on annotation of depressive symptoms before PD diagnosis (Premorbid depression, n = 9) versus those without annotation preceding PD diagnosis (Depression, n = 23) and compared both groups to the group without depression annotation (No Depression, n = 54). Both premorbid depression and depression groups showed increased eigengene values compared to the non-depressed group (Supplementary Figure 6). A pairwise comparison of the three groups with a Wilcoxon rank sum test, with BH correction found a significant difference between the non-depressed and depressed group (q-value = 0.02) whilst no significant difference was observed between any of the other groupwise comparisons with the premorbid or non-depressed group.

### Genes annotated from depression-associated DNA co-methylation in PD show ontological enrichment for synaptic processes

Next, to gain insight into potential underlying molecular functions captured by these trait-associated modules, we performed Gene Ontology (GO) analysis in the missMethyl package, a method which tests for enrichment for gene symbols annotated to each methylated loci whilst correcting for coverage bias of the 450K array^35^. The DepressionSN module showed nominal enrichment (P value <0.01) for 28 terms; within the top 10 most enriched terms (Figure 3, Supplementary Table 2), several were related to synaptic function, including synapse, maintenance of synapse structure and presynaptic active zone. Two additional terms were related to corticotropin releasing hormone. In addition, we also observed terms of extrinsic component of membrane, cellular component maintenance and sodium channel regulator activity. KEGG pathway enrichment identified 10 pathways with nominal enrichment (P value < 0.05), the topmost containing multiple pathways relevant to signalling (mTOR, Insulin, Phospholipase D, Apelin and Adrenergic) and longevity regulation (Supplementary Table 3).

**Figure 3:**
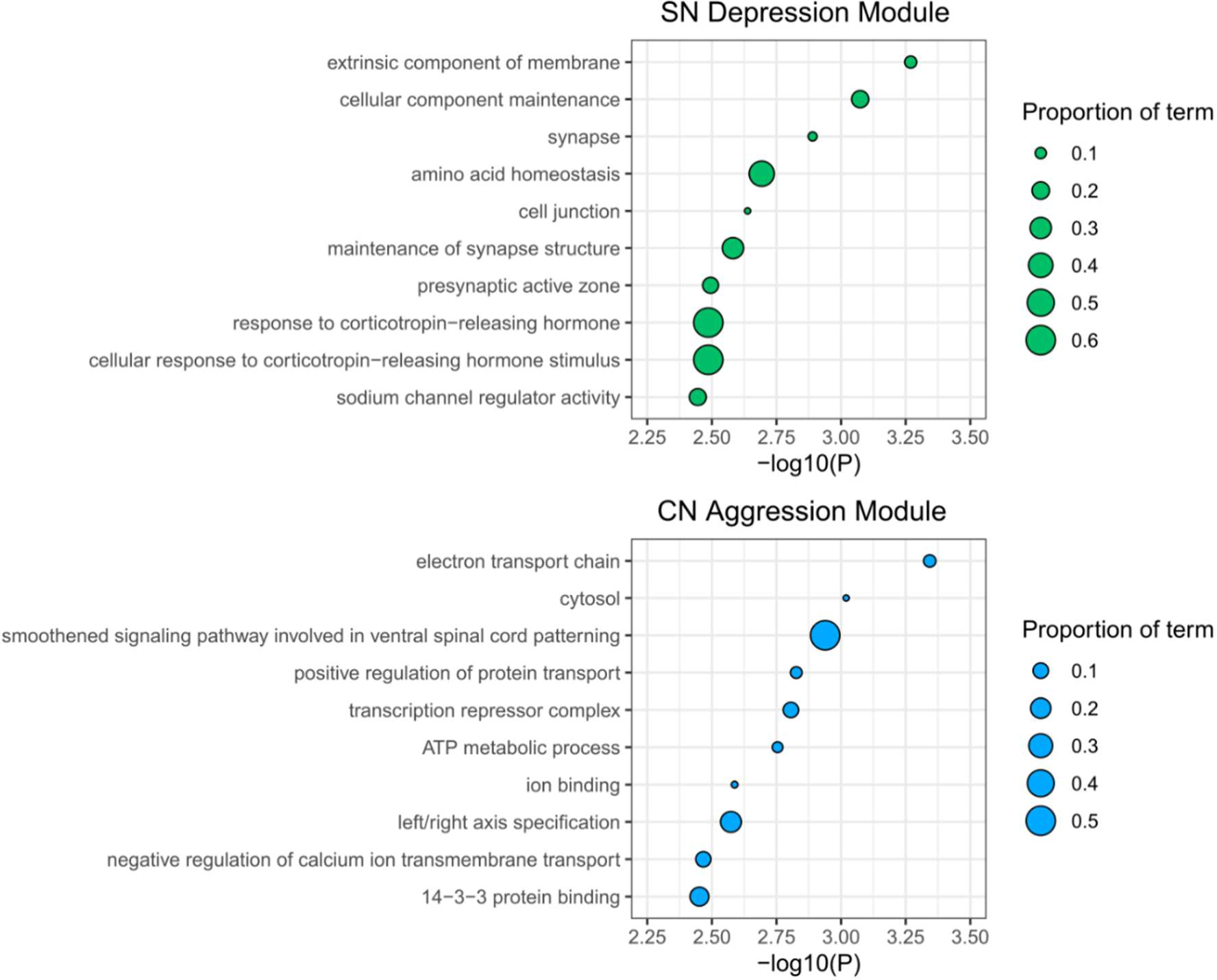
Gene Ontology Enrichment plots. The top 10 most significant terms for gene ontology (GO) enrichment analyses in **A)** the SN Depression associated module and **B)** the CN Aggression associated module. The term titles are displayed along the Y-axes, with -log10(P) for enrichment significance shown along the X-axis. Points are sized by the proportion of the overall ontology gene numbers represented in that specific module.

The AggressionCN module showed nominal enrichment (P value <0.01) for 49 terms, the top 10 most enriched (Figure 3, Supplementary Table 4), being related to electron transport chain, cytosol, spinal cord signalling, protein transport, transcription repression, ATP metabolism, ion binding, calcium ion transport and 14-3-3 protein binding. KEGG pathway enrichment identified 36 pathways with nominal enrichment (P value < 0.05) and included pathways relevant to carbon metabolism, diabetes and notch signalling (Supplementary Table 5).

### Genes annotated to the depression-associated DNA co-methylation module show significantly enriched expression in neuronal subtypes of the substantia nigra

As a number of terms in the gene ontology analysis of the DepressionSN module indicated neuronal involvement, we next sought to elucidate the cell-specific expression of genes annotated to the DNA methylation loci in the DepressionSN module, using a reference set of human single nucleus RNAseq data^36^ generated in the SN and using Expression Weighted Cell Enrichment analysis (EWCE) to test for enrichment. Of the 617 genes overlapping between the DepressionSN module methylation dataset and the reference snRNAseq data, we observed significant enrichment of expression in neurons only (Figure 4, Supplementary Table 6). Of a total of 68 defined cell subtypes annotated in the original study by Kamath and colleagues, 15 showed a significant expression enrichment for DepressionSN annotated genes (BH corrected q value < 0.05), corresponding to seven excitatory neuronal populations, four inhibitory neuronal populations and four dopaminergic neuronal populations. Of these populations, the two with the highest standard deviation shift from the mean expression were both excitatory: POSTN and OPRD1 (7.44 and 6.35 standard deviations from the mean, respectively). This provides evidence that the network of genes present within the DepressionSN module has functional relevance in disease as it is likely driven by SN neuronal cell types.

**Figure 4:**
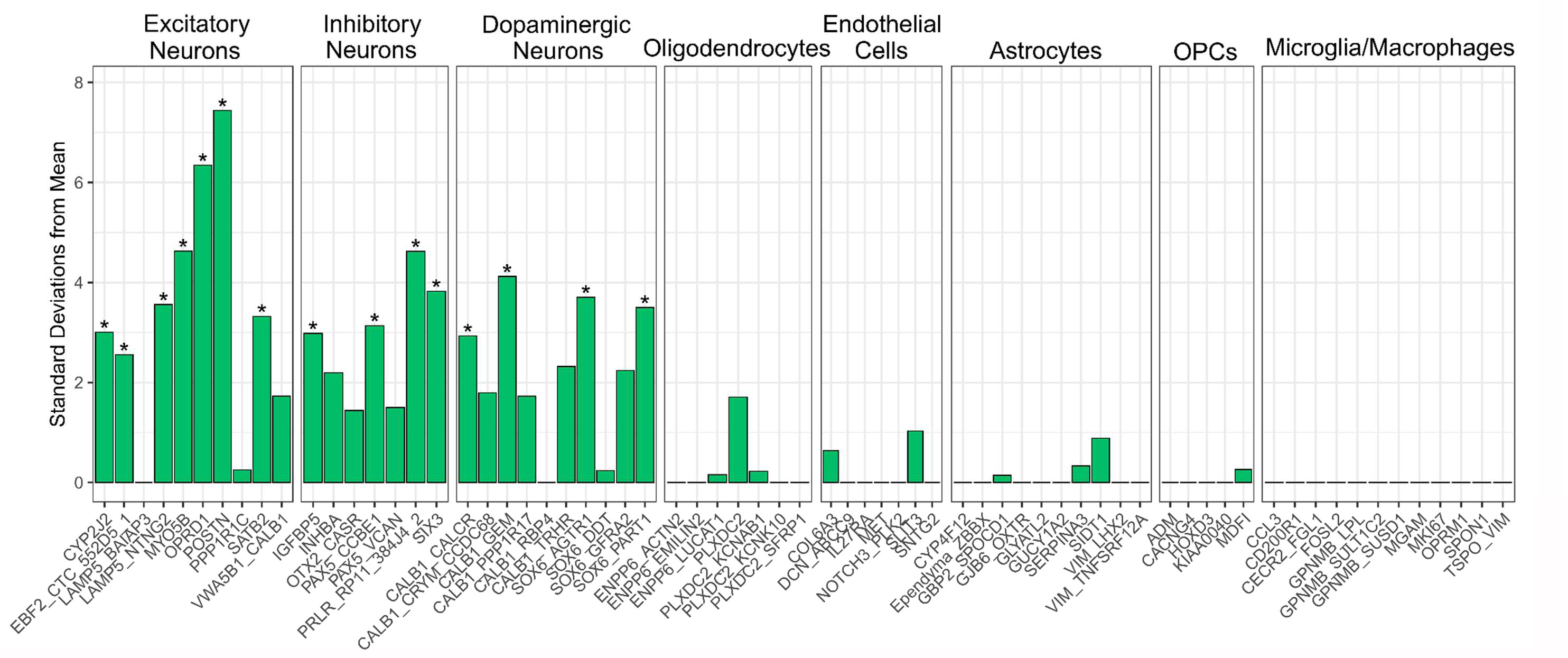
Expression Weighted Cell Enrichment analysis of DepressionSN module genes within the snRNAseq dataset from Kamath et al^36^. Sub-cell type annotations along the X-axis are grouped by their broader cell class. Benjamini-Hochberg (BH)-corrected significant enrichments (q < 0.05) are annotated with asterisks. Standard deviations from the mean value is displayed along the Y-axis. Standard deviation from the mean indicates the number of standard deviations from the mean level of expression of genes in the DepressionSN module, relative to the bootstrapped mean for that cell type.

## Discussion

In this report we have investigated multiple common secondary symptom traits in PD across three disease-relevant brain regions and explored the contribution of DNA methylation (summarized into inter-correlated DNA methylation networks) with trait presentation. We report region-specific associations between DNA methylation networks and trait presentation, specifically in association with depression in the SN and aggression in the CN. Subsequent downstream analyses indicated that genes related to the depression-associated SN co-methylation network (DepressionSN module) are enriched for ontological terms corresponding to synaptic processes, with significant overrepresentation of genes that are expressed in neuronal cells in the SN inferred from a separate snRNAseq dataset.

Depression in PD has a prevalence of roughly 40-46%^37^ and is a common premorbid symptom, being a risk factor for both PD development^14^ and worse symptom progression over time^38^. The pathophysiology underlying depressive symptoms in PD however remains poorly understood, with multiple potential threads of evidence for its etiology and relation to PD pathological development. Our results relating to SN neuronal changes lend support to dopaminergic theories of PD depression onset. Previous studies have shown that depressed PD patients present with greater neuronal loss^39^ and gliosis^40^ in the SN than non-depressed patients. Furthermore, alpha-synuclein pathology in the SN has been reported to be significantly higher in the SN of depressed cases versus non-depressed^39^. This regional neurodegeneration and consequent disruption of dopaminergic neurotransmission may be a contributing factor to the epigenetic alterations we observe in our results. However, the epigenetic network identified appears to be most enriched for expression in non-dopaminergic excitatory and inhibitory neurons, contradicting the evidence that this effect is purely a result of dopaminergic neurodegeneration. Further research is needed to fully elucidate the contribution of these SN neuronal cell types in the context of PD depression.

A potential avenue for further research could be in animal models of PD neurodegeneration, specifically in the context of PD depression. A number of rodent studies utilizing neurotoxic compounds such as 1-Methyl-4-Phenyl-1,2,3,6-Tetrahydropyridine (MPTP), which cause selective dopaminergic neuron degeneration, report depression-like behaviours, even manifesting before the onset of motor impairments^41^. Importantly the onset of this depression-like phenotype is variable^42, 43^, potentially allowing for a controlled model for assessment of specific cell type contributions to the variable onset of PD depression, as mediated by midbrain dopaminergic degeneration. Interestingly, in the original publication of the single nucleus RNA reference data used in our study, Kamath et al^36^ reported consistencies in dopaminergic subtypes between rodents and primate cell types, but reported one cell subtype characterized by expression of *CALB1* and *GEM* (“CALB1_GEM”) to be highly specific to primates. We observe this cell subtype as one of the four dopaminergic populations with significant enrichment of PD depression-associated network genes. This may have implications for the translatability of findings between rodent models and human disease in the context of PD depression.

As is a common issue with DNA methylation studies, in particular in bulk tissue, the causality of any changes detected is unclear, in particular in a disease process where cell type proportion changes are implicit. Although we have controlled for inferred cell type proportions, we cannot exclude the fact that the perturbed DNA methylation network we observe in the SN may be a downstream consequence of broad neurodegeneration in this brain region. Furthermore, it is premature to conclude whether differential DNA methylation of genes present within this particular network lead to altered expression, an assumption relied-upon for the findings of the snRNAseq enrichment analysis. Further work, in appropriate powered cohorts to look at a depression trait within PD and testing gene expression changes in the SN is required to validate this. However, our study represents the first of its type to look at underlying epigenomic changes with multiple symptom changes in PD and provides a basis for replication to confirm our findings.

A caveat to our findings is in the nature of the phenotyping data present and the clinical binary subsetting used in our trait annotations. Although care has been taken to annotate these records ante-mortem, we are limited in our ability to resolve clinical traits and inaccuracy may be present within this labelling criteria. In particular, we do not have capacity to resolve timing of symptoms for certain individuals and are limited to binary presentation over lifetime. This may have had a detrimental impact on identifying significant findings for specific outcomes tested in this study. Ideally, the use of quantitative scoring criteria, for example, the geriatric depression scale, Unified Parkinson’s Disease Rating Scale, or neuropsychiatric inventory (lacking in the case notes available for the current cohort) in future studies will allow for standardization of these traits.

## Conclusions

To conclude, we find evidence of regional epigenetic changes in relation to the development of secondary symptoms in PD, investigating multiple common secondary symptom traits in PD across three relevant brain regions and exploring the contribution of DNA methylation (summarised into inter correlated networks) with trait presentation. We find brain region-specific correlations between these networks and trait presentation, specifically in association with depression in the SN and highlight relevant ontological terms enriched within this network. Finally, we find that expression of genes within this network are specifically enriched for expression within relevant neuronal subtypes, prioritizing neuronal changes in the SN and cell types with potential contribution to the onset of PD depression.

## Methods

### Parkinson’s Sample Summary

PD samples profiled in this study have been summarized previously^44^. Tissue for 134 unique individuals was sourced from the Parkinson’s Disease UK Brain Bank, covering the Substantia Nigra (SN), Caudate Nucleus (CN) and Frontal Cortex (FC). Samples were excluded for having atypical parkinsonism noted on pathology reports, or early onset of disease (age of onset < 40). For our analysis, case notes were assessed for lifetime prevalence of secondary symptoms of depression, anxiety, aggression, dementia, hallucinations and sleep disturbance. As an example, for the annotation of hallucinations, evidence of the following psychiatric sections of clinical notes were used to evidence presence in three separate cases:

1. “deterioration, cognitive decline with visual hallucinations-animals”
2. “Hallucinations (visual, auditory, tactile?)
3. “Hallucinations; Confusion”

Whereas three examples with evidence of absence of hallucinations had the following psychiatric sections of the clinical notes:

1. “Nightmares - vivid dreams; Anxiety; Poor memory (late); No dementia, no hallucinations”
2. “Somnolence & lethargy; impaired memory; dementia”
3. “Poor concentration (losing train of thought)”

Where relevant, acute symptoms were not included (e.g. situational short term depression). For sensitivity analysis of premorbid depression, premorbid depression was evidenced either by explicit annotation in the clinical notes (e.g. “Anxiety and depression prior to onset of motor symptoms”) or from temporal staging of dated symptom entries. Years of disease, as defined by the number of years between diagnosis and death was also annotated as a separate outcome.

### DNA methylation profiling

Genome wide DNA methylation was profiled using the Illumina 450K methylation array which interrogates ∼450,000 methylation sites across the genome and has been described previously^45^. Data underwent stringent quality control and normalization as previously described using functions available in the *wateRmelon*^46^ R package (version 1.26). Samples with low median methylated or unmethylated intensities (n = 0) and with low bisulfite conversion percentages as determined using the *bscon()* function (n = 2) were removed as part of quality control (QC). Using a principal component (PC) based method, samples were tested for overlap of reported and predicted biological sex and removed if discordant (n = 2). Using single nucleotide polymorphism (SNP) probes included on the array, samples were checked for expected genetic relatedness for replicates across multiple brain regions (n = 12 removed for discordant expected relatedness). Using the *pfilter()* function samples were excluded if >1% of probes showed a detection value > 0.05 (n = 0) and probes were excluded if showing >1% of samples with detection value >0.05 or beadcount <3 in 5% of samples (n = 2,411 probes). Samples were tested for outliers using the *outlyx()* function and visually assessed using PC analysis. As a subtle separation of data points on the PC analysis could be seen corresponding to the different brain regions, we normalized each brain region separately. Quantile normalization was conducted using the *dasen()* function with default settings. Normalization violence was assessed using the *qual()* function to determine samples with high degrees of difference between raw and normalized beta values, with no outlying samples apparent.

### Weighted Gene Correlation Network Analysis

#### Data processing and module detection

Due to the high number of CpG sites tested on the Illumina 450K array and the low groupwise sample size available for this sample set, we aimed to reduce the multiple testing burden for association using Weighted Gene Correlation Network Analysis (WGCNA, version 1.69)^47^. WGCNA determines correlation networks present within a given dataset to identify distinct clusters of highly correlated data-points which may hold functional relevance based on its significant pairwise relationship to other datapoints. Datasets were first filtered for variable probes across each brain region separately as determined by median absolute deviation (MAD) for any individual probe > median MAD for the entire dataset. To ensure consistent probes were being fed into each analysis per brain region, all probes passing this threshold in every brain region were included for further analysis, resulting in a set of 243,783 probes. Following this, all brain regions were processed separately. To reduce the effect of unwanted technical and biological variance, multiple regression was used to regress out these effects from the dataset. Each CpG site was regressed against age, sex, technical batch, NeuN+ predicted cell proportion (estimated using the *estimateCellCounts()* function in waterMelon^48^) and post mortem interval (PMI). The residuals from this regression were extracted and added to the intercept to give a methylation value controlling for the applied covariates and scaled similarly to the raw value. Residual corrected methylation values were then clustered by Euclidean distance and the first four PCs were visually assessed to test for outlying samples. From this analysis two samples were removed from the SN, one from the FC and one from the CN. Co-methylation network and module detection were determined in a block-wise method and set as unsigned, so to weight correlation between probes irrespective of the direction of correlation. As recommended in the WGCNA protocol, soft thresholding was applied, which raises the power of each correlation to a particular value with the aim to reduce noise within the dataset. A scale free topology graph was constructed for powers ranging from 1-20 in stepwise increments and assessed for balance between scale free topology and connectivity. From this, a value of eight was selected for the SN, 12 for the FC and nine for the CN. Finally, modules were identified using the *blockwiseModules()* function (unsigned network, min size = 100, max size = 10000).

#### Module Trait Association Analysis

Identified modules ranged in size and similarity and were labelled based on an arbitrary color value determined by the WGCNA process. Modules were additionally filtered at this point to remove any remaining modules retaining any significant (P <0.05) confounding trait association. For association testing, CpGs present within each module were aggregated into individual values representative of a weighted average of methylation within that module. These values, termed module eigengenes (MEs) are calculated using the eigenvalues from the first PC for all methylation values in that modules with one module eigengene value determined for each individual case in the dataset per module. These module eigengenes were tested for association with phenotypic traits, using Pearson’s correlation for continuous traits and Spearman’s correlation for binary traits. For multiple testing correction, Bonferroni correction was applied as 0.05 divided by the number of modules present in that particular brain region.

#### Module membership analysis

For modules that showed a significant association with any of the outcomes tested, individual relevant probes were assessed based on module membership (MM) and probe significance (PS). MM is calculated using Pearson’s correlation between an individual probe and the ME of the module it is assigned and is thus representative of that individual probe’s connectivity to the rest of the module. PS was determined using correlation analysis between individual methylation values and the trait found to be significantly associated with that modules eigengene in the same method as for overall ME association. MM was tested against –log10 transformed PS using Pearson‘s correlation.

#### Ontological enrichment analysis

For ontological enrichment analysis, annotated gene-symbols from the Illumina manifest file were extracted for the corresponding modules. We used a background of all annotated gene-symbols for all 243,783 probes fed into the analysis. Modules were tested for ontological terms for biological pathways enriched in CpGs present within each module using Gene Ontology (GO) and Kyoto Encyclopedia of Genes and Genomes (KEGG) analysis using the missMethyl package^35^ (version 1.30). As similar ontology terms were observed from this output due to overlapping gene sets, modules were merged based on semantic similarity using the web tool REVIGO (http://revigo.irb.hr/)^49^. Resnik’s measure was used to compute the similarity of terms and a medium between terms similarity of 0.7 was allowed.

### Single cell data processing and cell enrichment analysis

To determine the sub cellular localization of annotated genes determined from the WGCNA analysis, human SN single nuclei RNA sequencing (snRNAseq) data generated using the 10X Genomics (v.3) kit for the Kamath et al. 2022^36^ publication was sourced from the Broad Institute Single Cell Portal https://singlecell.broadinstitute.org/single_cell/study/SCP1768/. Filtered human single nuclei barcodes, gene features and expression matrix along with processed UMAPs and metadata were downloaded and processed. Data was loaded using the *Read10X()* function in the *Seurat* R package (version 4.3.0). Loaded data was converted to a summarized experiment object using the *SummarizedExperiment()* function in the R package of the same name. colData for each profiled cell was assigned from the corresponding celltypes based on annotations from the UMAP files as determined from the original publication and annotated at two levels of granularity. From this, a set of 37,389 genes were taken forward. Data was then processed for Expression Weighted Cell Type Enrichment analysis using functions within the *EWCE*^50^ package (https://github.com/NathanSkene/EWCE) (version 1.4.0). First, genes with no overall expression (n = 4236 genes) or no significant differential expression between cell types (n = 3,532 genes at FDR adjusted q-value threshold <1e-05) were removed using the *drop_uninformative_genes()* function with the Limma setting. A normalized mean expression and specificity cell type dataset was calculated using the *generate_celltype_data()* function. Data quality was assessed at this point for potential artefacts by visual assessment of known marker gene expression in known cell types using the *plot_ctd()* function. Genes annotated to methylated loci in each module determined from WGCNA were tested separately for cell type enrichment using the *bootstrap_enrichment_test()* function. Tests were conducted over 100,000 repetitions and tested for cell type and sub-cell type enrichment separately (Supplementary Figure 7). Significant cell type enrichment was determined by Benjamini-Hochberg (BH) corrected q-values < 0.05. For plotting, similar modules were determined based on Euclidean distance of a binary significance module by cell type matrix.

## Author contributions

LH and KB conducted laboratory experiments generating the DNA methylation data. JH undertook data analysis and bioinformatics, with support from RGS, IC and EP. ARS, LSW and BC collated and interpreted the clinical data for the analysis. NW and KL conceived the project. EP and KL supervised the project. JH, EP and KL drafted the manuscript. All authors read and approved the final submission.

## Acknowledgments

This work was funded by research grants from the Medical Research Council (MRC (MR/S011625/1), the National Institute of Aging (NIA) of the National Institutes of Health (NIH) (R01AG067015) and BRACE to KL and from Parkinson’s UK (Project Grants G-1309 and G-1502) to NW. This study was supported by the National Institute for Health and Care Research Exeter Biomedical Research Centre. The views expressed are those of the authors and not necessarily those of the NIHR or the Department of Health and Social Care.

## Competing interests

The authors declare no competing interests.

## Code availability

All codes are available at https://github.com/JoshHarveyGit/PD_TraitNetworkAnalysis

## Notes

### Competing Interest Statement

The authors have declared no competing interest.

## References

1. Ray Dorsey, E., et al. Global, regional, and national burden of Parkinson’s disease, 1990–2016: a systematic analysis for the Global Burden of Disease Study 2016. Lancet Neurol. 17, 939–953 (2018).

2. Jankovic, J. Parkinson’s disease: clinical features and diagnosis. J. Neurol. Neurosurg. Psychiatry 79, 368–76 (2008).

3. Jankovic, J. & Tan, E. K. Parkinson’s disease: etiopathogenesis and treatment. J. Neurol. Neurosurg. Psychiatry 91, 795–808 (2020).

4. Institute, K. et al. Depression in Parkinson disease—epidemiology, mechanisms and management. Nat. Rev. Neurol. 2011 81 8, 35–47 (2011).

5. Broen, M. P. G., Narayen, N. E., Kuijf, M. L., Dissanayaka, N. N. W. & Leentjens, A. F. G. Prevalence of anxiety in Parkinson’s disease: A systematic review and meta-analysis. Mov. Disord. 31, 1125–1133 (2016).

6. Aarsland, D. et al. Range of neuropsychiatric disturbances in patients withParkinson’s disease. J. Neurol. Neurosurg. Psychiatry 67, 492 (1999).

7. den Brok, M. G. H. E., et al. Apathy in Parkinson’s disease: A systematic review and meta-analysis. Mov. Disord. 30, 759–769 (2015).

8. Aarsland, D. et al. Parkinson disease-associated cognitive impairment. Nat. Rev. Dis. Prim. 2021 71 7, 1–21 (2021).

9. Aarsland, D. et al. Parkinson disease-associated cognitive impairment. Nat. Rev. Dis. Prim. 2021 71 7, 1–21 (2021).

10. Jones, S., Torsney, K. M., Scourfield, L., Berryman, K. & Henderson, E. J. Neuropsychiatric symptoms in Parkinson’s disease: aetiology, diagnosis and treatment. BJPsych Adv. 26, 333–342 (2020).

11. Aarsland, D., Larsen, J. P., Tandberg, E. & Laake, K. Predictors of nursing home placement in Parkinson’s disease: a population-based, prospective study. J. Am. Geriatr. Soc. 48, 938–942 (2000).

12. Fénelon, G. & Alves, G. Epidemiology of psychosis in Parkinson’s disease. J. Neurol. Sci. 289, 12–17 (2010).

13. Munhoz, R. P. et al. Demographic and motor features associated with the occurrence of neuropsychiatric and sleep complications of Parkinson’s disease. J. Neurol. Neurosurg. Psychiatry 84, 883–887 (2013).

14. Bareeqa, S. B. et al. Prodromal depression and subsequent risk of developing Parkinson’s disease: a systematic review with meta-analysis. https://doi.org/10.2217/nmt-2022-0001 12, 155–164 (2022).

15. Gustafsson, H., Nordström, A. & Nordström, P. Depression and subsequent risk of Parkinson disease. Neurology 84, 2422–2429 (2015).

16. Schuurman, A. G. et al. Increased risk of Parkinson’s disease after depression: a retrospective cohort study. Neurology 58, 1501–1504 (2002).

17. Sandor, C. et al. Universal clinical Parkinson’s disease axes identify a major influence of neuroinflammation. Genome Med. 14, 1–15 (2022).

18. Greenland, J. C., Williams-Gray, C. H. & Barker, R. A. The clinical heterogeneity of Parkinson’s disease and its therapeutic implications. Eur. J. Neurosci. 49, 328–338 (2019).

19. Liu, G. et al. Genome-wide survival study identifies a novel synaptic locus and polygenic score for cognitive progression in Parkinson’s disease. Nat. Genet. 2021 536 53, 787–793 (2021).

20. Creese, B. et al. Glucocerebrosidase mutations and neuropsychiatric phenotypes in Parkinson’s disease and Lewy body dementias: Review and meta-analyses. Am. J. Med. Genet. Part B Neuropsychiatr. Genet. 177, 232–241 (2018).

21. Allis, C. D. & Jenuwein, T. The molecular hallmarks of epigenetic control. Nat. Rev. Genet. 2016 178 17, 487–500 (2016).

22. MacBean, L. F., Smith, A. R. & Lunnon, K. Exploring Beyond the DNA Sequence: A Review of Epigenomic Studies of DNA and Histone Modifications in Dementia. Curr. Genet. Med. Reports 2020 83 8, 79–92 (2020).

23. Roubroeks, J. A. Y. et al. An epigenome-wide association study of Alzheimer’s disease blood highlights robust DNA hypermethylation in the HOXB6 gene. Neurobiol. Aging 95, 26–45 (2020).

24. Lunnon, K. et al. Methylomic profiling implicates cortical deregulation of ANK1 in Alzheimer’s disease. Nat. Neurosci. 2014 179 17, 1164–1170 (2014).

25. Smith, R. G. et al. A meta-analysis of epigenome-wide association studies in Alzheimer’s disease highlights novel differentially methylated loci across cortex. Nat. Commun. 2021 121 12, 1–13 (2021).

26. Shao, X. et al. Dementia with Lewy bodies post-mortem brains reveal differentially methylated CpG sites with biomarker potential. Commun. Biol. 2022 51 5, 1–11 (2022).

27. Chuang, Y. H. et al. Longitudinal Epigenome-Wide Methylation Study of Cognitive Decline and Motor Progression in Parkinson’s Disease. J. Parkinsons. Dis. 9, 389–400 (2019).

28. Young, J. I. et al. Genome-wide brain DNA methylation analysis suggests epigenetic reprogramming in Parkinson disease. Neurol. Genet. 5, e342 (2019).

29. Pihlstrøm, L. et al. Epigenome-wide association study of human frontal cortex identifies differential methylation in Lewy body pathology. Nat. Commun. 2022 131 13, 1–10 (2022).

30. Pishva, E. et al. Psychosis-associated DNA methylomic variation in Alzheimer’s disease cortex. Neurobiol. Aging 89, 83–88 (2020).

31. Chuang, Y.-H. et al. Longitudinal Epigenome-Wide Methylation Study of Cognitive Decline and Motor Progression in Parkinson’s Disease. J. Parkinsons. Dis. 9, 389–400 (2019).

32. Braak, H., Alafuzoff, I., Arzberger, T., Kretzschmar, H. & Tredici, K. Staging of Alzheimer disease-associated neurofibrillary pathology using paraffin sections and immunocytochemistry. Acta Neuropathol. 112, 389–404 (2006).

33. Braak, H., Ghebremedhin, E., Rüb, U., Bratzke, H. & Del Tredici, K. Stages in the development of Parkinson’s disease-related pathology. Cell Tissue Res. 318, 121–134 (2004).

34. Aarsland, D. et al. Parkinson disease-associated cognitive impairment. Nat. Rev. Dis. Prim. 2021 71 7, 1–21 (2021).

35. Wang, Z., Wu, X. L. & Wang, Y. A framework for analyzing DNA methylation data from Illumina Infinium HumanMethylation450 BeadChip. BMC Bioinformatics 19, 15–22 (2018).

36. Kamath, T. et al. Single-cell genomic profiling of human dopamine neurons identifies a population that selectively degenerates in Parkinson’s disease. Nat. Neurosci. 2022 255 25, 588–595 (2022).

37. Jellinger, K. A. The pathobiological basis of depression in Parkinson disease: challenges and outlooks. J. Neural Transm. 2022 12912 129, 1397–1418 (2022).

38. Ng, A., Chander, R. J., Tan, L. C. S. & Kandiah, N. Influence of depression in mild Parkinson’s disease on longitudinal motor and cognitive function. Park. Relat. Disord. 21, 1056–1060 (2015).

39. Patterson, L., Rushton, S. P., Attems, J., Thomas, A. J. & Morris, C. M. Degeneration of dopaminergic circuitry influences depressive symptoms in Lewy body disorders. Brain Pathol. 29, 544–557 (2019).

40. Frisina, P. G., Haroutunian, V. & Libow, L. S. The neuropathological basis for depression in Parkinson’s disease. Parkinsonism Relat. Disord. 15, 144 (2009).

41. Mou, Y. K. et al. Application of Neurotoxin-Induced Animal Models in the Study of Parkinson’s Disease-Related Depression: Profile and Proposal. Front. Aging Neurosci. 14, 504 (2022).

42. Zhang, X. et al. Decrease of gene expression of astrocytic 5-HT2B receptors parallels development of depressive phenotype in a mouse model of Parkinson’s disease. Front. Cell. Neurosci. 9, (2015).

43. Tang, J. et al. Crocin Reverses Depression-Like Behavior in Parkinson Disease Mice via VTA-mPFC Pathway. Mol. Neurobiol. 57, 3158–3170 (2020).

44. Kia, D. A. et al. Identification of Candidate Parkinson Disease Genes by Integrating Genome-Wide Association Study, Expression, and Epigenetic Data Sets. JAMA Neurol. 78, 1 (2021).

45. Kia, D. A. et al. Identification of Candidate Parkinson Disease Genes by Integrating Genome-Wide Association Study, Expression, and Epigenetic Data Sets. JAMA Neurol. 78, 1 (2021).

46. Pidsley, R. et al. A data-driven approach to preprocessing Illumina 450K methylation array data. BMC Genomics 14, 293 (2013).

47. Langfelder, P. & Horvath, S. WGCNA: An R package for weighted correlation network analysis. BMC Bioinformatics 9, 1–13 (2008).

48. Pidsley, R. et al. A data-driven approach to preprocessing Illumina 450K methylation array data. BMC Genomics 14, 1–10 (2013).

49. Supek, F., Bošnjak, M., Škunca, N. & Šmuc, T. REVIGO Summarizes and Visualizes Long Lists of Gene Ontology Terms. PLoS One 6, e21800 (2011).

50. Skene, N. G. & Grant, S. G. N. Identification of vulnerable cell types in major brain disorders using single cell transcriptomes and expression weighted cell type enrichment. Front. Neurosci. 10, 16 (2016).

