## Supplementary figures for "Epigenetic insights into neuropsychiatric and cognitive symptoms in Parkinson’s disease: A DNA co-methylation network analysis"

**Supplementary Figure 1: Symptom Overlap Matrices**. Grids display in correlation matrix format the proportion of overlap in paired symptom presentation. **A)** shows this as a percentage overlap, interpreted as the percentage of samples with symptoms displayed along the Y-axis that also have the symptom along the X-axis. For example, the top left value of 47.92% refers to the percentage of samples with sleep disturbance that also had dementia. **B)** shows the same information but depicted as raw total sample numbers with particular symptom overlap.

A)

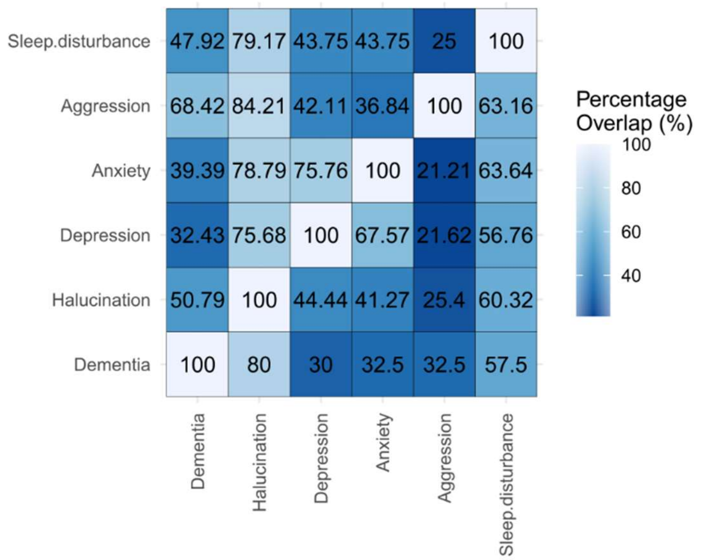


B)

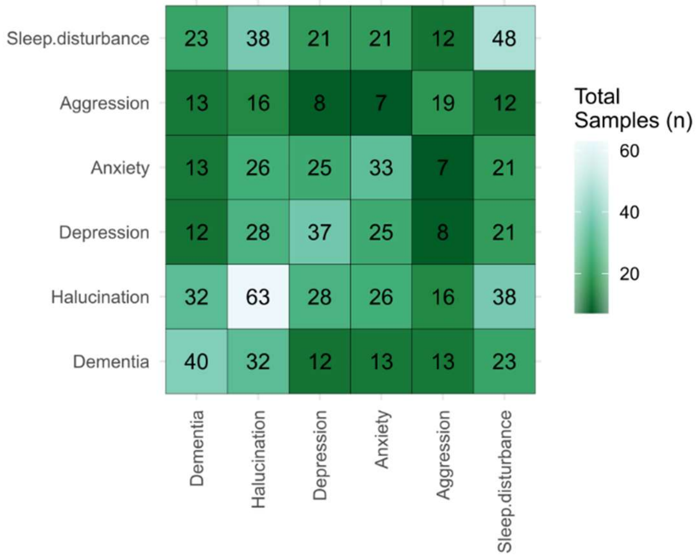

**Supplementary Figure 2: Trait-module correlation matrices for the substantia nigra.** Correlations are shown between module eigengenes and traits of interest, with module names (arbitrarily assigned colors) shown along the Y-axis. Correlation estimates are reported, with p-values in parentheses. Grids are colored by correlation estimates. Abbreviations: PMI: Post mortem interval, NeuN_pos: Predicted NeuN+ proportion, batch: Processing batch value, AgeAtDeath: Years of age at death, YrsDisease: Years between diagnosis of PD and death.

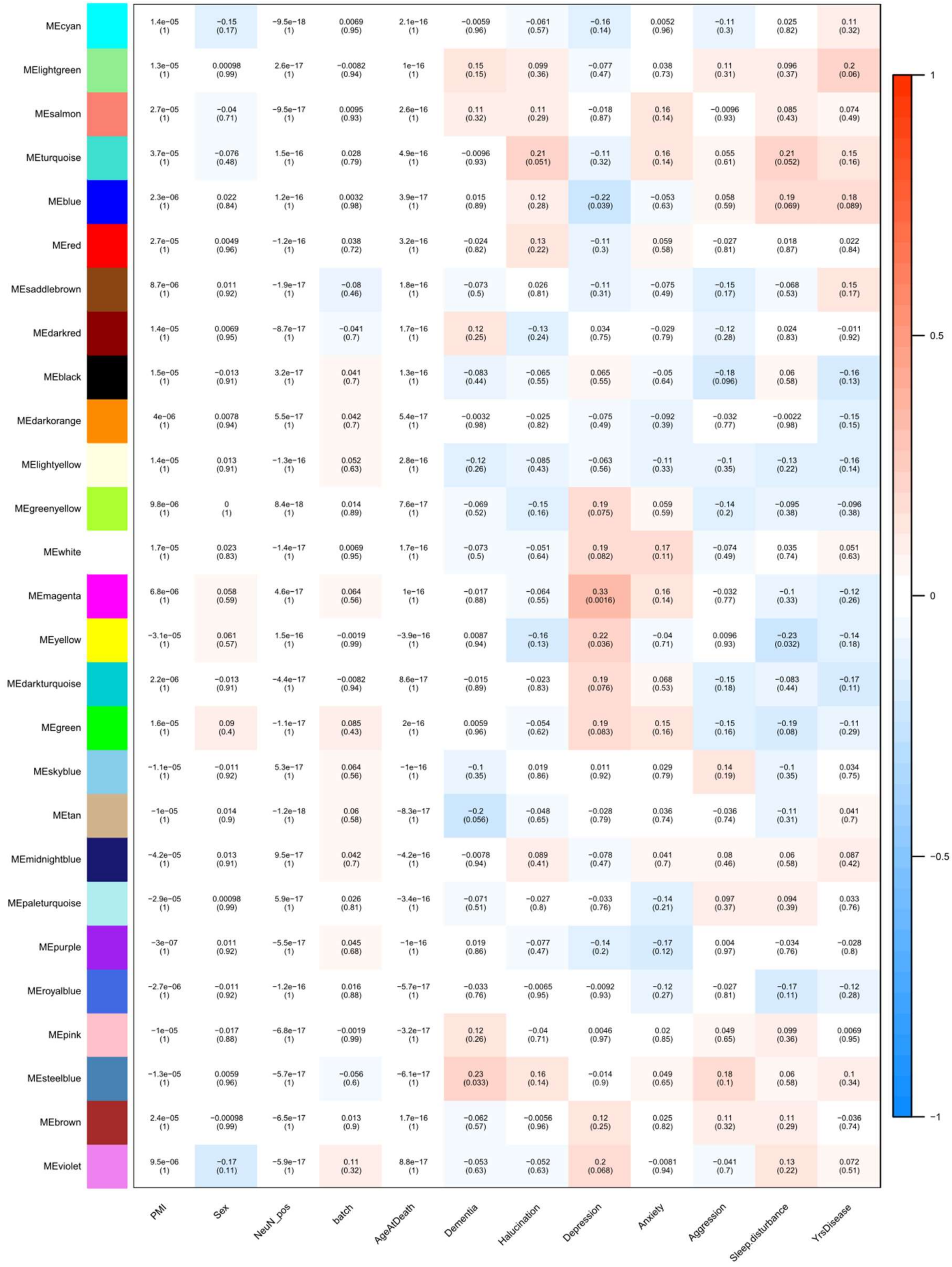

**Supplementary Figure 3: Trait-module correlation matrices for the frontal cortex.** Correlations are shown between module eigengenes and traits of interest, with module names (arbitrarily assigned colors) shown along the Y-axis. Correlation estimates are reported, with p-values in parentheses. Grids are colored by correlation estimates. Abbreviations: PMI: Post mortem interval, NeuN_pos: Predicted NeuN+ proportion, batch: Processing batch value, AgeAtDeath: Years of age at death, YrsDisease: Years between diagnosis of PD and death.

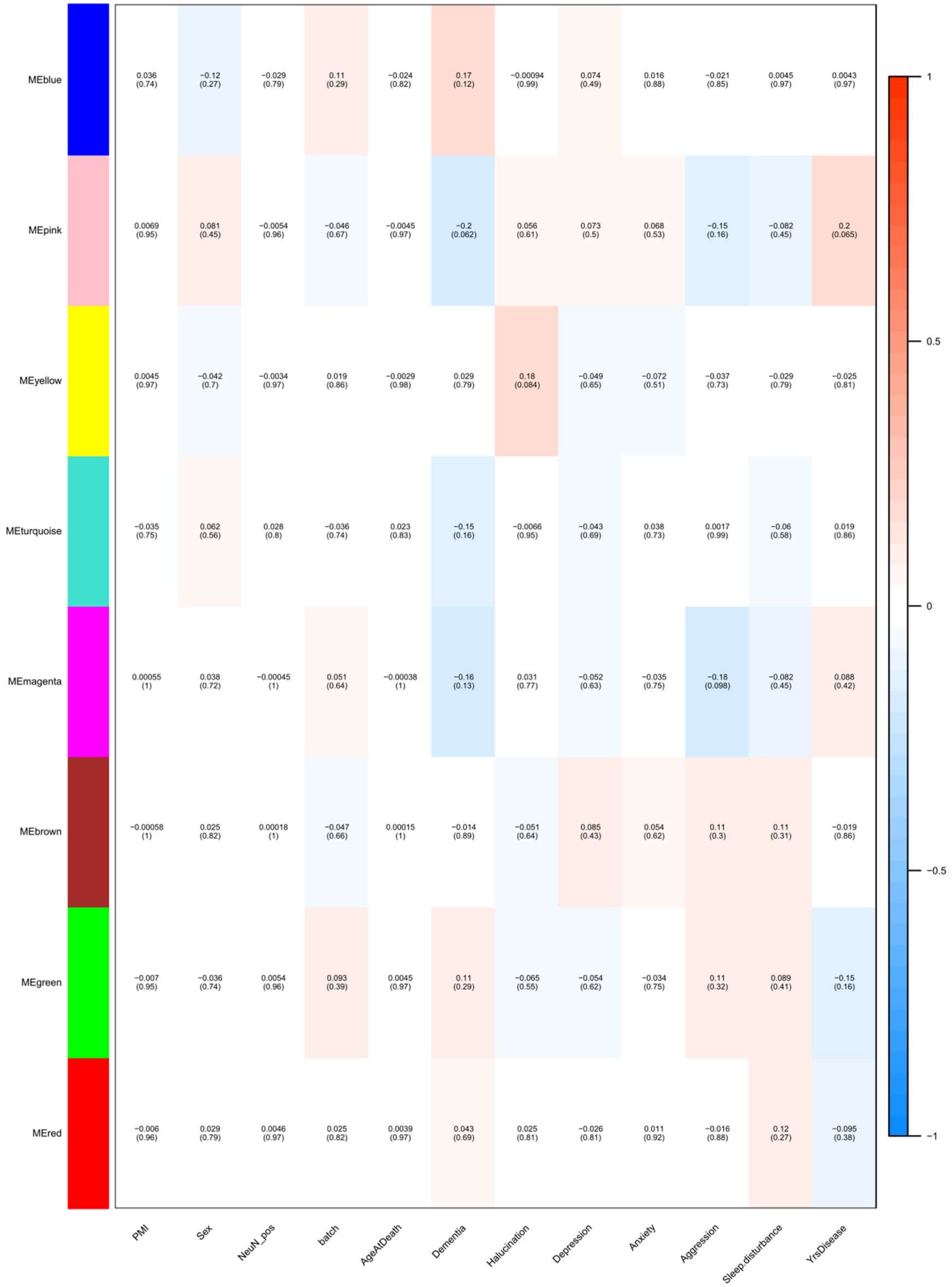

**Supplementary Figure 4: Trait-module correlation matrices for the caudate nucleus.** Correlations are shown between module eigengenes and traits of interest, with module names (arbitrarily assigned colors) shown along the y-axis. Correlation estimates are reported, with p-values in parentheses. Grids are colored by correlation estimates. Abbreviations: PMI: Post mortem interval, NeuN_pos: Predicted NeuN+ proportion, batch: Processing batch value, AgeAtDeath: Years of age at death, YrsDisease: Years between diagnosis of PD and death.

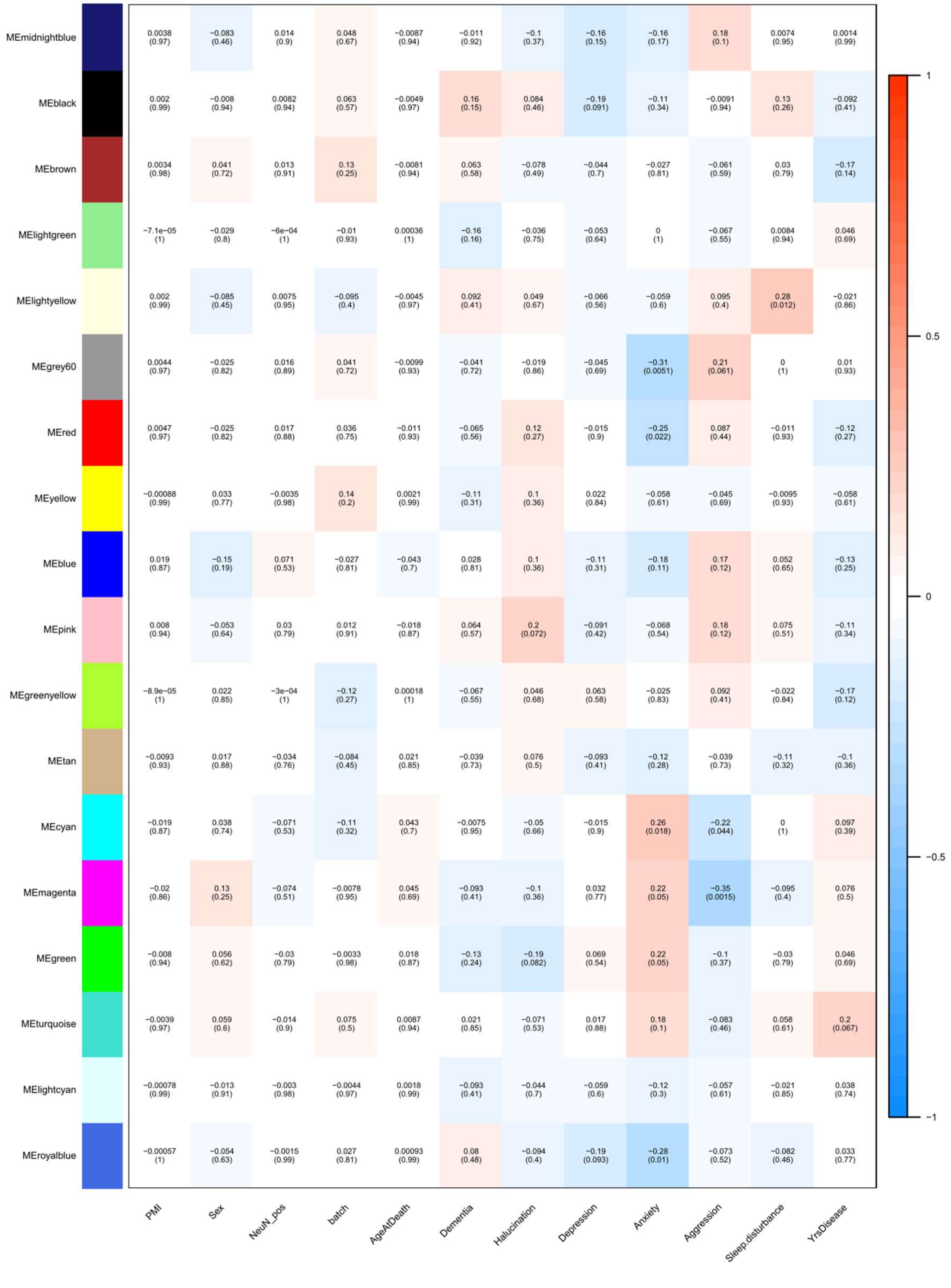

**Supplementary Figure 5: Correlation between module membership and gene significance for trait tested.** Module membership (here measured as the correlation coefficient between the individual probes corrected methylation value and the module eigengene), and the gene significance (-log10 transformed p-value) from Spearman's correlation between each probe and the trait being tested. Measurements are calculated for all probes within the **A)** DepressionSN module (n = 1,375 probes, Pearson's Correlation Coefficient = 0.12, p-value = 1.24e-05) and **B)** AggressionCN module (n = 475 probes, Pearson's Correlation Coefficient = 0.07, p-value = 0.13).

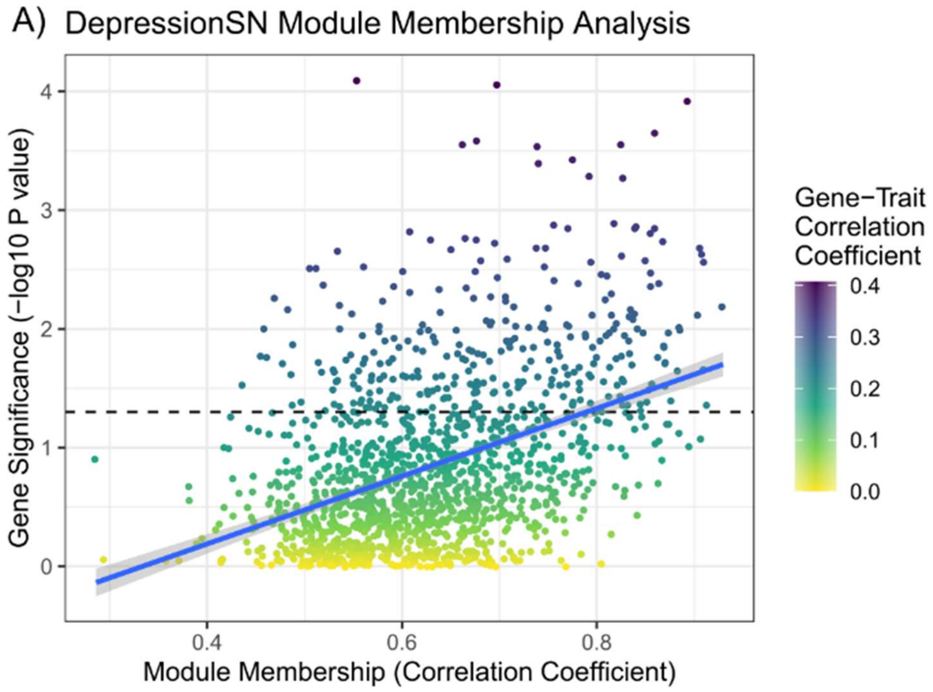


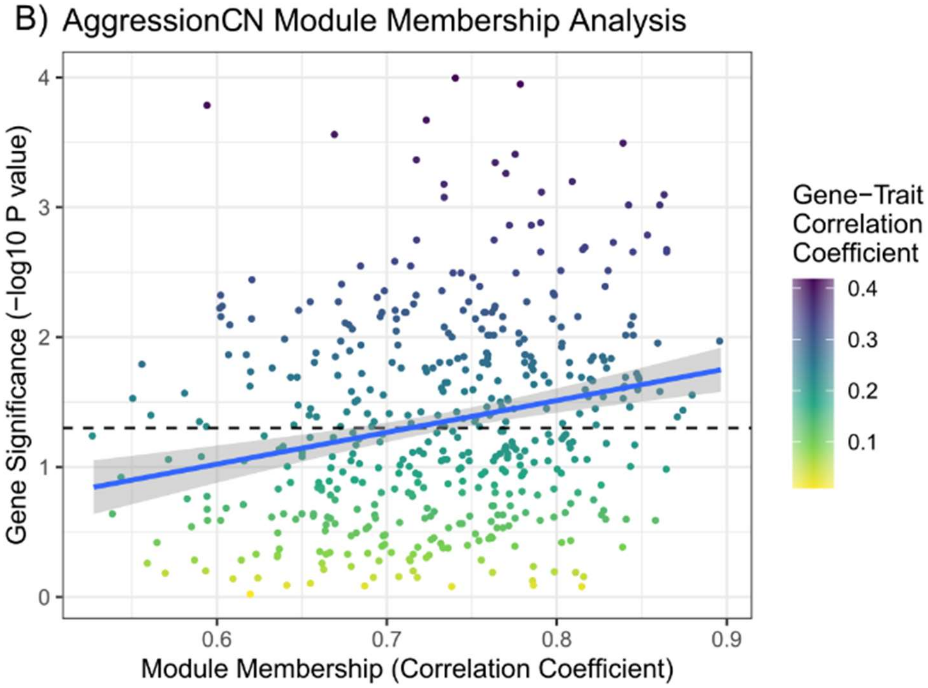

**Supplementary Figure 6: Violin and boxplots of depressionSN module eigengene compared to depression onset.** Groups are subset by temporal annotation of depression symptoms in the clinical notes. Premorbid depression is a history of depression preceding the primary PD diagnosis. Depression is a group with no annotated history of depression before the primary PD diagnosis. Significance annotated for a pairwise Wilcox rank-sum comparison between each group with BH-correction. * indicates q-value < 0.05

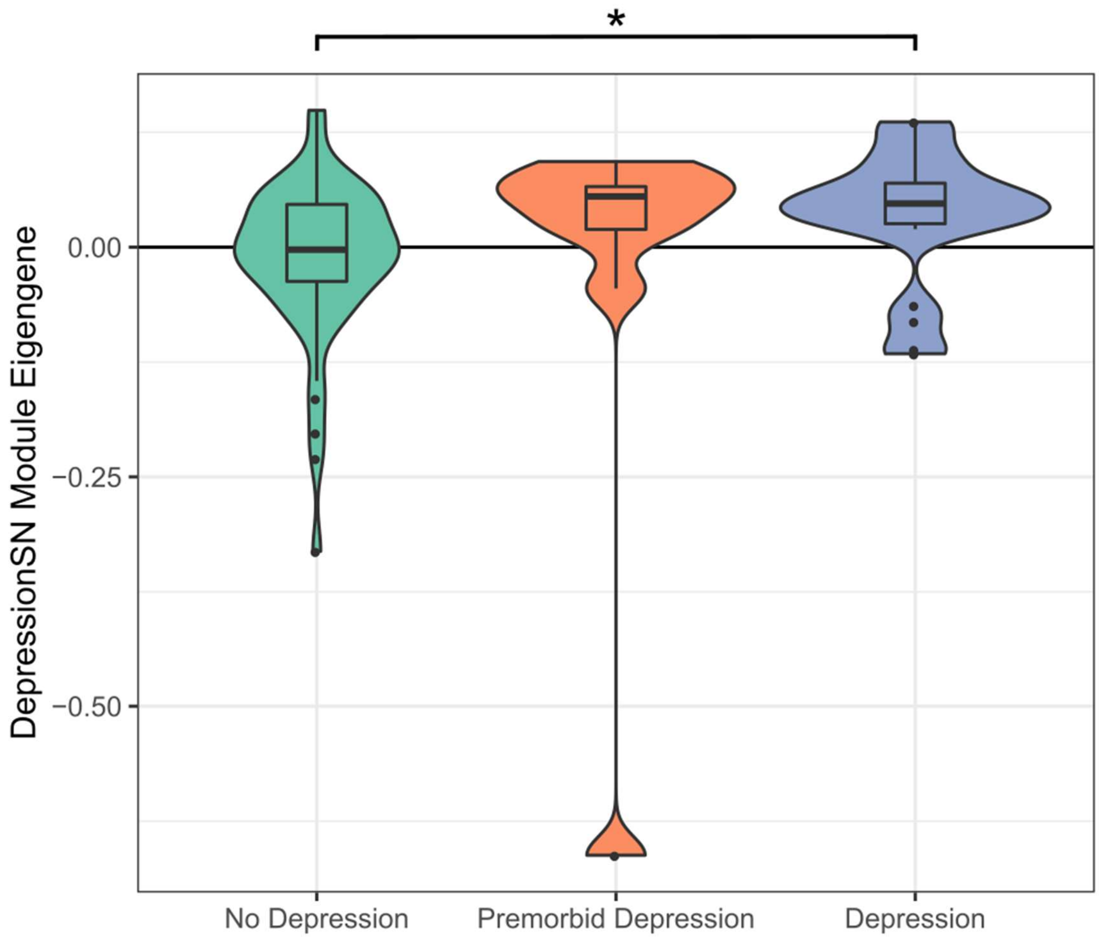

**Supplementary Figure 7: Expression Weighted Cell Type Enrichment Results for all modules detected in the substantia nigra.** Results are displayed as a binary matrix format, with cells coloured based on Benjamini-Hochberg (BH) significant correct p-value for significant enrichment correcting for all 252 separate tests. Modules are shown along the Y-axis and cell types tested for significant enrichment are shown along the X-axis. Highlighted with a black box is the magenta module, corresponding to the DepressionSN module

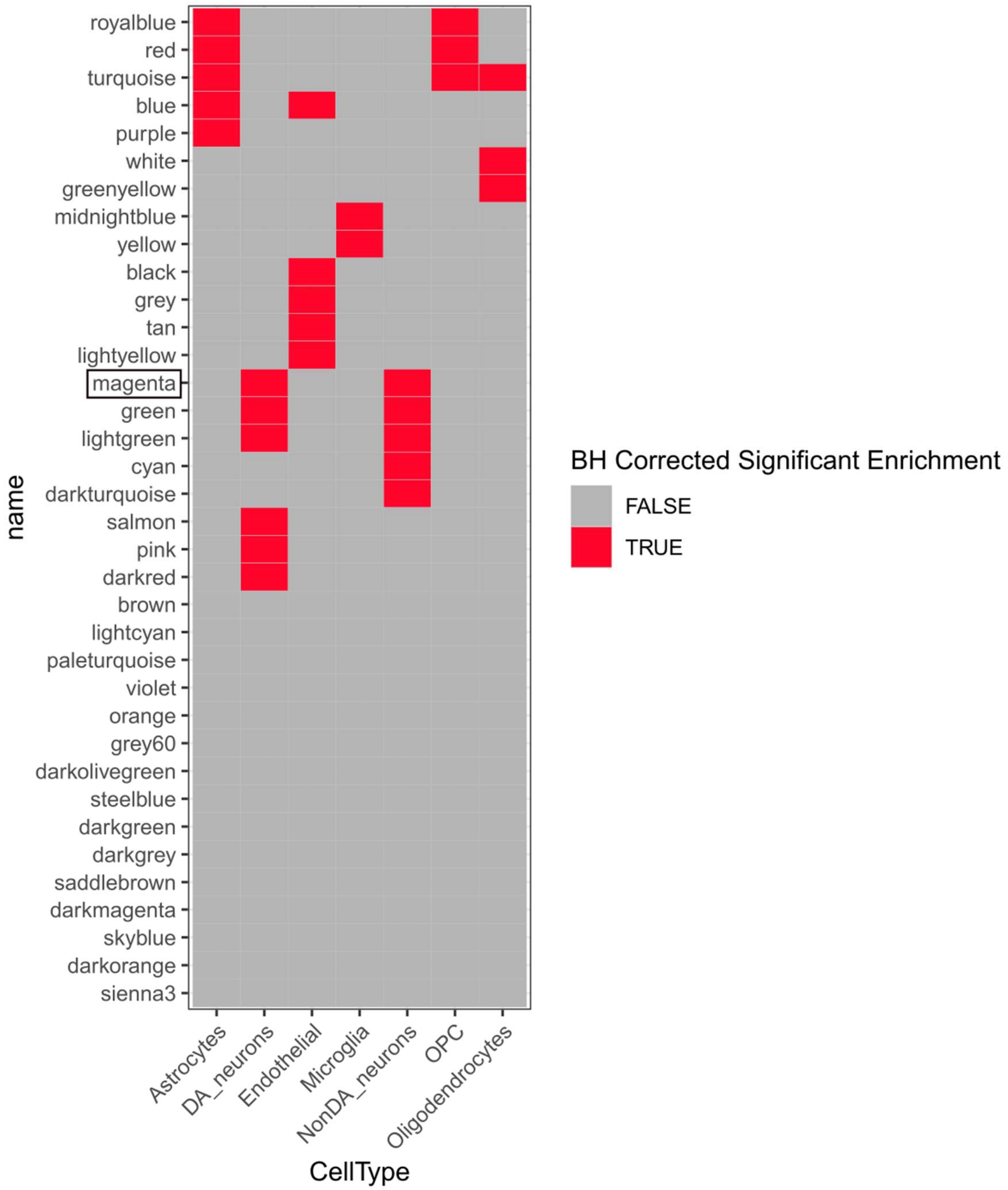
